# Lead in the head: persistent lead poisoning of waterfowl in the Camargue (southern France) ten years after the ban on the use of lead shot in wetlands

**DOI:** 10.1101/2024.01.29.577719

**Authors:** Arnaud Béchet, Anthony Olivier, François Cavallo, Lou Sauvajon, Jocelyn Champagnon, Pierre Defos du Rau, Jean-Yves Mondain-Monval

## Abstract

Lead pollution remains a worldwide environmental and health issue with persistent detrimental effects on human and wildlife. Despite having been identified as an issue for wildlife a long time ago, the use of lead shot for hunting in wetlands was banned only in 2006 in France. Here, we took advantage of a long-term monitoring of waterfowl lead shot contamination in the Camargue (southern France) to (1) assess the local effectiveness of the French regulation in reducing waterfowl contamination and to (2) assess local compliance regarding the use of nontoxic shots in wetlands from 2007 onwards. We collected waterfowl gizzard from 38 hunters in the Camargue during 20 hunting seasons (1998 to 2017) for a total of 2306 gizzards from 28 different species. From 2008 to 2019, we also systematically collected the cartridge casings at three sites of communal wetland hunting. Ratio of lead vs nontoxic cartridges found in the field allowed monitoring hunter compliance with the new regulation. Despite a ten-year long ban on lead shot our results do not show any significant reduction in waterfowl exposure to lead through shot ingestion in the Camargue over the 20-year monitoring period. Indeed, gizzards of harvested waterfowl were as likely to contain at least one lead shot before the ban as after with a mean prevalence of 12% over the 13 species considered across the study period. Cartridge casings showed the persistence of use of lead shots by hunters in the Camargue wetlands despite a slow increase in the use of nontoxic cartridges. This unequivocally indicates that the law is far from being strictly applied, pointing to an insufficient policy enforcement. Our results support the need for a complete ban of lead ammunition for both wetland and terrestrial wildfowl which will facilitate policy enforcement and compliance.

## Introduction

Lead pollution remains a worldwide environmental and health issue with persistent detrimental effects onto humans and wildlife (Pain et al. 2019; Larsen and Sánchez-Triana 2023). Several public policies have been implemented at national and international scales to reduce the risks of lead contamination (Avery 2009; Lacerda et al. 2023). Lead hunting ammunition remains one of the largest sources of lead discharged into the environment in Europe, and it places many wildlife taxa at risk of being poisoned (Treu et al. 2020). In particular, a long lasting issue has been the contamination of waterfowl by spent hunting lead shots ingested as grit while foraging in wetlands (Fig. 1). Despite having been identified as an issue for wildlife a long time ago (Bowles 1908; Wetmore 1919; Bellrose 1959; Hoffmann 1960), it took a large amount of scientific evidence and long lasting efforts of advocacy before policy regulations aiming at removing this specific threat were eventually implemented (Arnemo et al. 2016; Treu et al. 2020).

**Figure 1.**
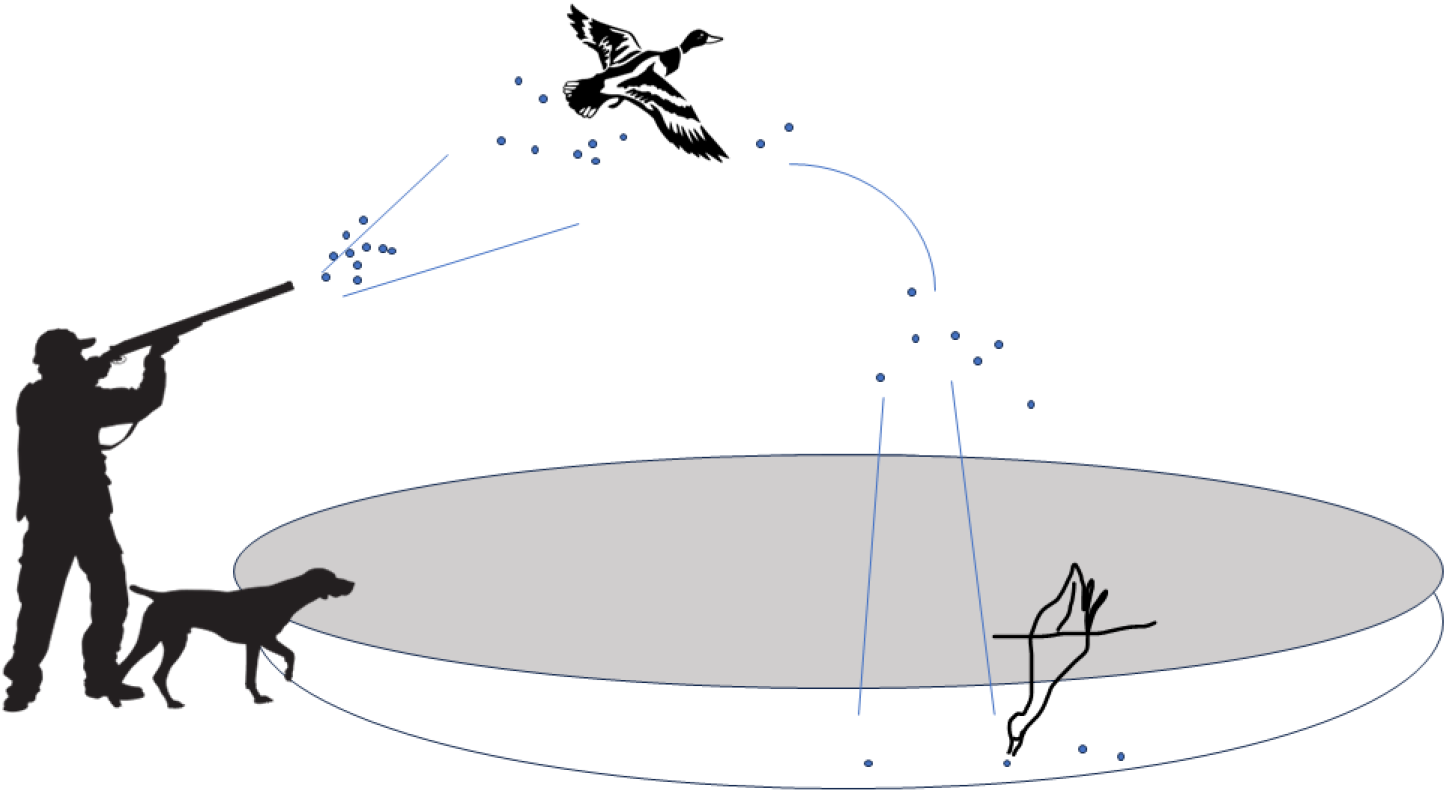
Process by which spent gunshot are consumed as grit by waterfowl.

In Europe, the use of lead shots was first forbidden in 1986 in Denmark Ramsar wetlands (Mateo and Kanstrup 2019). Later, many countries introduced legislation in response to the African-Eurasian Waterfowl Agreement (AEWA) recommendation for the use of non-toxic shot over wetlands (Avery 2009). Implementation of the regulation on lead shot use for waterfowl remains a lengthy complicated process, since hunters in some European union countries are apparently complying well, e.g. Spain (Mateo et al. 2012) and Denmark (Kanstrup and Balsby 2019), while hunters from other countries are not, e.g in the UK (Stroud et al. 2021).

In France, the use of lead shot for hunting was banned in 2006 in wetlands but not for terrestrial zones. Such as forested areas, agricultural land, etc. As a result, this regulation proved difficult to enforce because it remained permitted (even in wetlands) to carry lead shot during a hunting session, as long as it was not used in wetlands. An overall review of this national policy was therefore necessary but, to our knowledge, there has been no evaluation of the effectiveness of this regulation on waterfowl contamination (but see Mondain-Monval et al. (2015) for the Camargue) nor of hunter compliance at national scale in France (Mondain-Monval et al. 2020).

The Camargue is France’s largest wetland (145,000 ha) and a major migration and wintering area for waterbirds in Europe (Galewski and Devictor 2016). Classified as a RAMSAR site, it is home to tens of thousands of Anatidae every winter (Tamisier and Dehorter 1999). The Camargue has a long tradition of waterfowl hunting, with several thousand waterfowl hunters harvesting waterfowl in the delta (Mondain-Monval et al. 2009). Such major historical activity has led to significant lead contamination of the marshes (Pain 1992) and of waterfowl alike (Hoffmann 1960; Hovette 1973; Pirot and Taris 1987; Pain 1990; Mondain-Monval et al. 2002, 2015).

In this study, we took advantage of a long-term monitoring of waterfowl lead shot contamination in the Camargue (Mondain-Monval et al. 2002) to (1) assess the local effectiveness of the French regulation in reducing waterfowl contamination, and (2) assess local compliance regarding the use of nontoxic shots in wetlands from 2007 onward.

## Material and methods

Waterfowl contamination is generally assessed by examining bird gizzards to quantify the number of ingested toxic *vs*. non-toxic pellets (Mondain-Monval et al. 2015). Bird gizzards were collected opportunistically from a network of hunters in the Camargue, during each hunting season from 1998/1999 to 2017/2018 (Fig. 2). A total of 38 different hunters contributed to the sampling effort. Those hunters harvested wildfowl both in communal hunting grounds and in private hunting estates. The number of gizzards collected per winter varied from 3 to 305 with a mean of 115 ± 20 (SE) gizzards collected per winter. A total of 2,306 gizzard were collected over the study period from 28 different species (Appendix 1).

**Figure 2.**
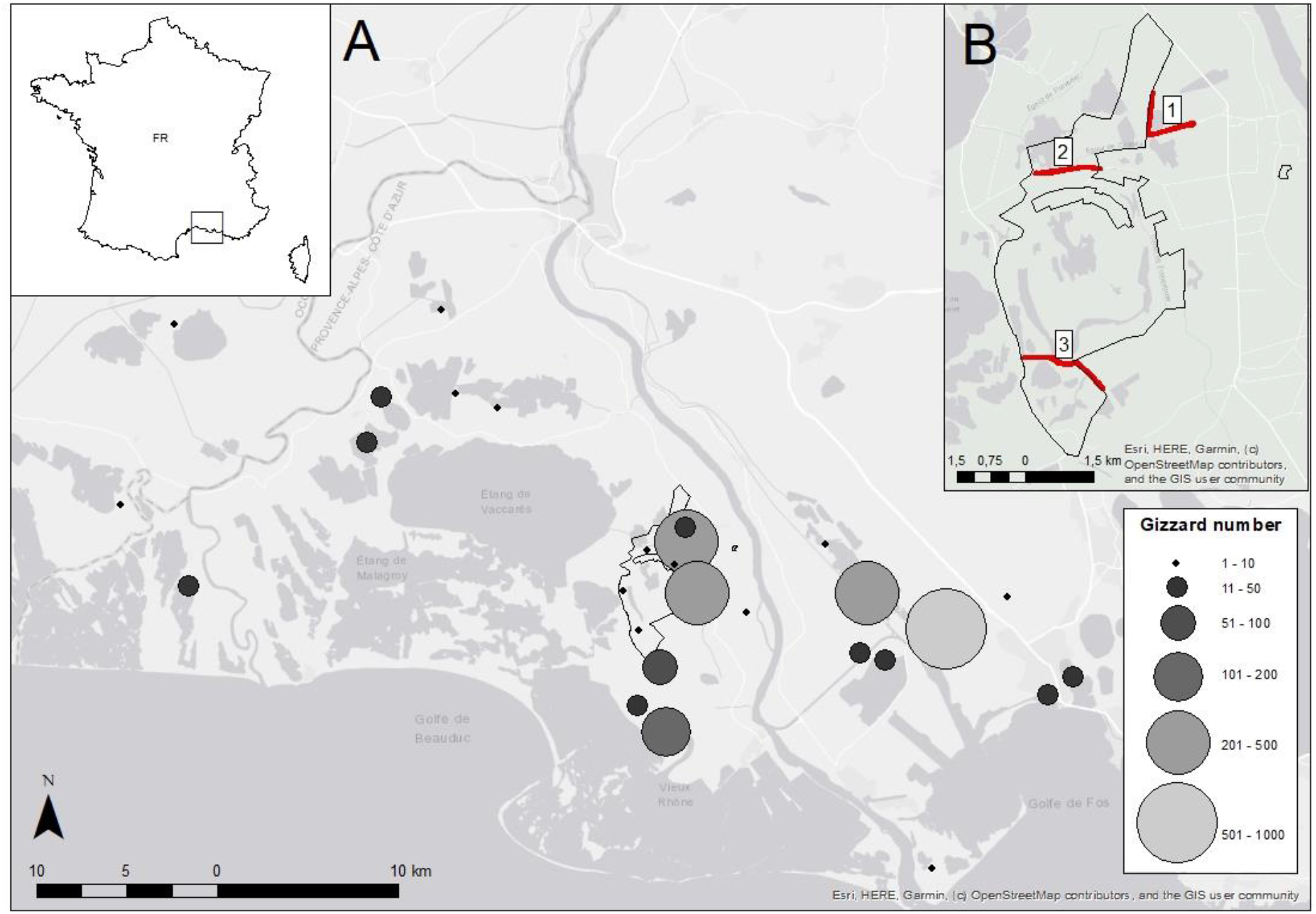
A) Origin of the gizzards collected by hunters from hunting season 1998/1999 to 2017/2018 in the Camargue. Circle size indicates the number of gizzards collected at each location. B) location of the communal hunting trails where cartridge casings were collected in the vicinity of the Tour du Valat nature reserve : 1) Draille du Sambuc, 2) Bordure du Fumemorte and 3) Draille Marseillaise

For each collected gizzard we recorded the species, date and exact location of harvest. Gizzards were deep frozen for later analysis. Two successive batches of gizzard examination were performed (in 2012 and in 2018). Gizzards were first examined visually to detect the presence of wounds. Indeed, unlike lead pellets, whether a steel pellet found in the gizzard was embedded (having penetrated through the gizzard flesh when the bird was shot) or ingested (swallowed by the bird as grit) is difficult to ascertain, as embedded steel pellets keep their original round shape and do not flatten on impact as lead pellets do. Steel pellets found in the gizzard were therefore considered to have been ingested if they had clearly been found eroded inside the gizzard, and embedded if the pellets showed no signs of wear or when wounding channels were present in the gizzard. Second, gizzards were cut in half and the contents washed into a dish with a water stream. Plant material and other food items were removed by flotation, and carefully inspected for any grit (including shot), which was retained in the dish. The remaining gizzard contents were examined by x-ray photographs to ensure that all shots were identified (Montalbano and Hines 1978) as the risk of missing some of them by the sole visual examination was high (especially when a gizzard contained a large number of lead shots or small eroded shots). Shot pellets attracted by a magnet were considered to be steel shot. In order to identify lead shot and the alloy composition of other shot types, all those not attracted by a magnet were analyzed by X-ray fluorescence spectrometer (Thermo Scientific XL3t 980 GOLDD +, Niton Corp.), with a 3 mm X-beam, metal alloys calibration mode and Kapton support.

In addition, every hunting season from 2008/2009 to 2019/2020, cartridge casings were collected at three public wetland hunting sites located on the outskirts of Tour du Valat nature reserve (Fig. 2; 43°30’30’’ N, 4°39’59’’ E). The cartridge casings abandoned by hunters on the ground were collected by walking several times during each hunting season along the 3 km of communal hunting paths (Draille du Sambuc, bordure du Fumemorte and Draille Marseillaise; Fig. 2) along which the communal hunters of Arles hunt (Groupe Cynégétique Arlésien). Only recent cartridge cases (no rusty marks) were considered in the analysis. Lead and nontoxic cartridges could be easily distinguished from the markings on the cartridge cases, so that hunter compliance to the regulation could be monitored (Mondain-Monval et al. 2020).

All hunting materials were collected in full accordance with French and European legislation, including the 2009/147/EC Directive, and Human and Animal Rights were fully respected in the framework of this study.

### Statistical analyses

#### Gizzard dataset

Because terrestrial species (i.e. European turtle dove *Streptopelia turtur*, Common pheasant *Phasianus colchicus* and Common quail *Coturnix coturnix*) were not the initial target of the lead ban, we removed them from the analyses. We only retained the game species with more than five gizzards collected in the dataset, so that species specific contamination rates be meaningful. Because we aimed at assessing the effect of lead ban on lead contamination, we only retained species for which there was at least one lead shot occurrence in them (Table 1). Hence, gizzards of Garganey *Spatula querquedula*, Tufted duck *Aythya fuligula*, Red knot *Calidris canutus*, Black tailed godwit *Limosa limosa*, Red-crested pochard *Netta rufina*, Eurasian Curlew *Numenius arquata*, Whimbrel *Numenius phaeopus*, Eurasian golden plover *Pluvialis apricaria*, Common greenshank *Tringa nebularia*, Redshank *Tringa totanus* and Northern lapwing *Vanellus vanellus* were discarded, so that the final dataset retains 13 species (but see Appendix 1 for a complete overview of shots found over the 28 species initially collected, including landbirds).

**Table 1.**
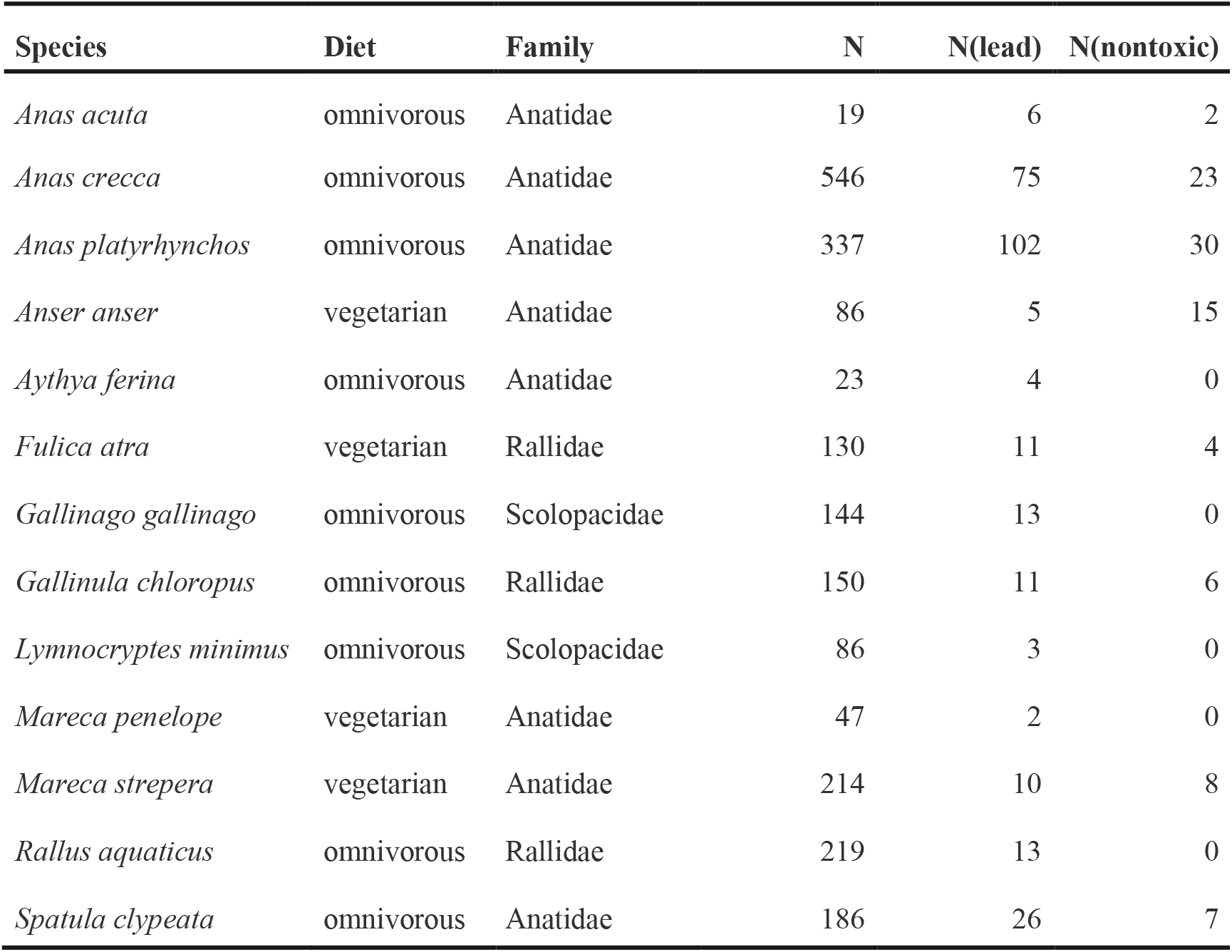
Species retained for the analysis of the effect of the lead ban on gizzard content in waterbirds of the Camargue (1998/99-2017/18) with their diet, family, number of Gizzard (N), number of gizzard containing at least one lead N(lead) and at least one nontoxic shot N(nontoxic).

Because lead toxicity starts with the first ingested lead shot (Tavecchia et al. 2001), we used the prevalence (presence/absence) of lead shot in gizzard as the response variable (number of ingested lead shot ranged from 0 to 104, with an average of 3.97 ± 0.55 lead shots per gizzard in the 281 birds with at least one ingested lead shot). Similarly, to assess the dynamics of replacement of lead by non-toxic shot as grit used by birds, we modeled the prevalence of non-toxic shot found in each gizzard. Finally, to control for variations in prevalence of shots ingested as grit that may be caused by species traits (i.e. diet or taxonomic family), we used the later factors as covariates in the models (Table 1). Species diet was determined following Cramps et al. (1988) with four categories (granivorous, carnivorous, omnivorous, and herbivorous; Appendix 1) for the initial dataset (28 species) was reduced to two levels (omnivorous or herbivorous) for the reduced dataset (13 species) that was further analyzed.

First, we modeled the variations of the probability of a gizzard to contain at least one lead shot using linear mixed models (logit link and binomial error term) with fixed effects being the ban period (before or after season 2006/2007), year, species functional traits (diet and family), and the random effect being species identity. The general model included the interactions between ban and diet, and between ban and family to assess possible differences in the effect of ban caused by species functional traits.

As the effect of the ban on lead shot prevalence may be slow and gradual, either because of poor hunter compliance with the new policy or because of lead shot persistence in wetlands (Kanstrup et al. 2020), we tested for a negative trend in the prevalence and number of lead shots in the gizzard from winter 2007 onward. We used linear mixed models with year, species trait and family as fixed effects and the species identity as a random effect. We modeled lead shot prevalence trend from winter 2007 onward using a binomial error distribution and lead shot numbers trend using a negative binomial error distribution. We also tested for alternative time trend shape of lead shot prevalence by using GAMM models with the exact same structure as the linear mixed model (hence keeping diet and family as covariables) and applying a smooth function on year.

Similarly, we modeled the variations of the probability of the gizzards containing at least one nontoxic shot using linear mixed models with fixed effects being the ban period, year, species trait and the random effect being species identity. Like for the lead analysis, we also evaluated if there was any trend in non-toxic shot prevalence in gizzard both from 2007 onward and over the whole study period.

We ran models with all combinations of effects which could be derived from the general model using the dredge function in R, and we ranked models using the Akaike information criterion (AICc; Burnham & Anderson (2002)). Then, we eventually averaged parameters of models with a delta AICc smaller than two. We discussed the effect of variables when the 95% confidence intervals of their estimate did not include 0. Linear mixed models were run using package glmmTMB of R (Magnusson et al. 2020). GAMM models were run using package mgcv of R (Wood 2001). Model averaging was performed using packages MuMin and function dredge (Bartón 2015).

#### Cartridge casings dataset

We assessed the effect of lead ban on the ratio of lead versus non-toxic cartridge found at hunting spots by running a simple logistic model with a binomial error term testing the effect of year on the probability of finding a lead cartridge. The response variable was computed as the number of lead cartridge over the total number of cartridges collected a given winter.

## Results

### Gizzard dataset

The reduced dataset resulted in 2187 gizzards from 13 species (see Table 1). 281 gizzards contained one or more lead shots, 96 contained one or more nontoxic shots. Fluorescence analysis allowed to detect one bismuth shot in a gizzard also containing both lead and steel shots and 14 gizzards with shots made of an alloy. In 12 of these alloy shots we found lead so they were in fact “improved lead shots”. All alloy shots were found in gizzards containing at least one lead shot hence we did not specifically analyzed the change in alloy shots across time (Appendix 2).

Model selection indicated that lead shot prevalence did not change from before to after the ban in bird gizzards (Fig. 3). The only explanatory variables retained in the best model (Table 2) were the covariates diet and family. There was a significant higher prevalence in omnivorous species than in vegetarian ones (β = -1.81 ± 0.55), and a significant lower prevalence in Rallidae than in the other families (β = -1.49 ± 0.49; Table 2). Overall, mean lead shot prevalence was 0.12 ± 0.007 but it varied a lot among species [range: 0.03-0.31] with *Anas acuta* and *Anas platyrhynchos* likely driving the diet effect as omnivorous species with 0.31 ± 0.10 and 0.30 ± 0.02 lead shot prevalence, respectively (Fig. 4).

**Table 2.** Model selection with degrees of freedom (df), AICc and ΔAICc, for the variation of the prevalence (probability of counting at least one shot) of lead shots ingested in the gizzard of the 13 species hunted in the Camargue (1998/99-2017/18) before and after the ban. Only models within 4 points of AIC with the best model are presented among the 13 models built.

| Model | df | AICc | ΔAICc |
| --- | --- | --- | --- |
| Intercept + diet + family | 5 | 1585.0 | 0 |
| Intercept + diet | 3 | 1587.1 | 2.10 |
| Intercept | 2 | 1588.7 | 3.69 |
| Intercept + family | 4 | 1588.7 | 3.70 |
| Intercept + ban + diet + family | 6 | 1588.9 | 3.91 |

**Figure 3.**
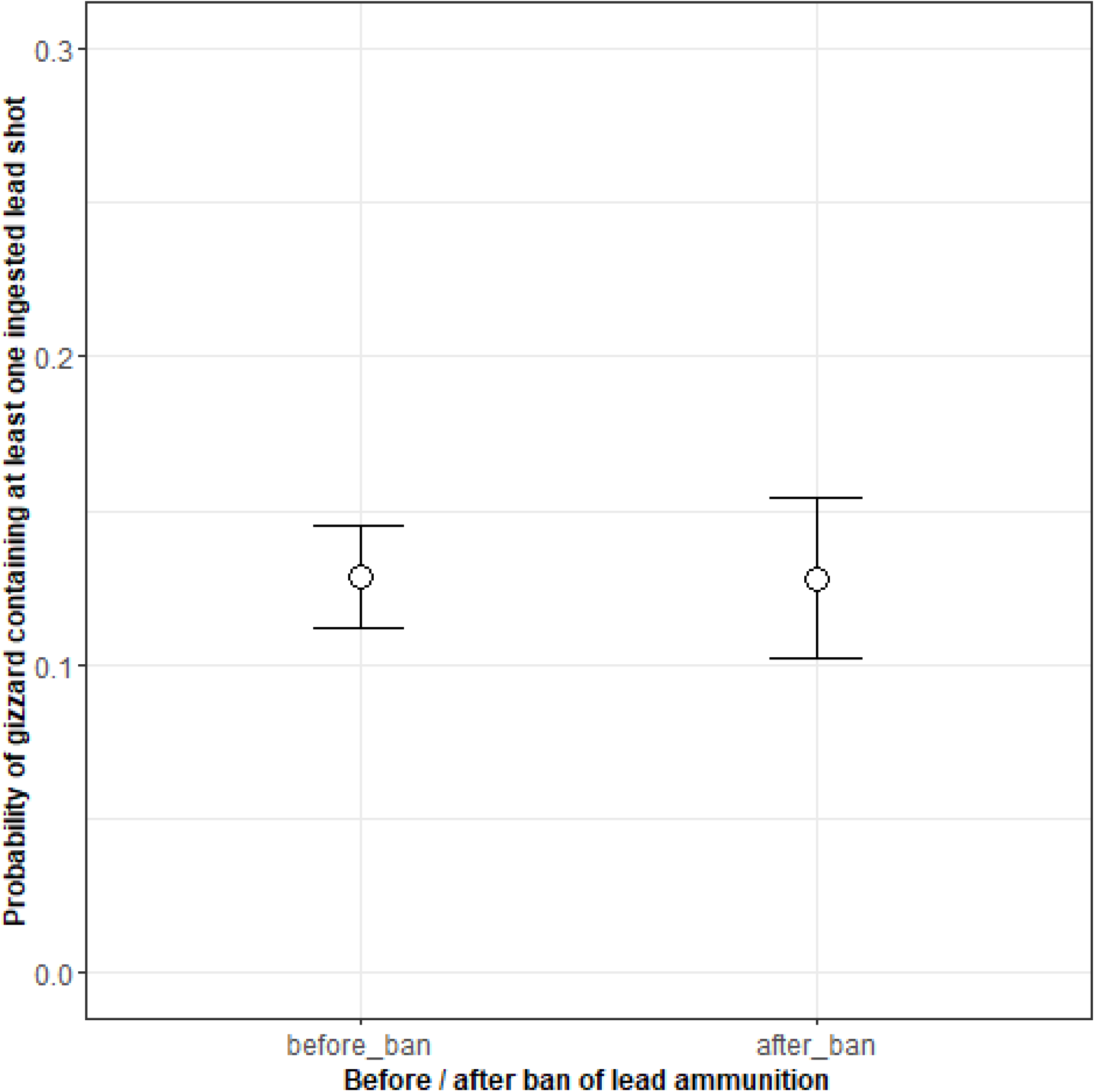
Prevalence of lead shot in the gizzards of the 13 waterbird species hunted in the Camargue (south of France) from hunting season 1998/1999 to 2017/2018 compared before (n= 1563) and after the ban (n=624).

**Figure 4.**
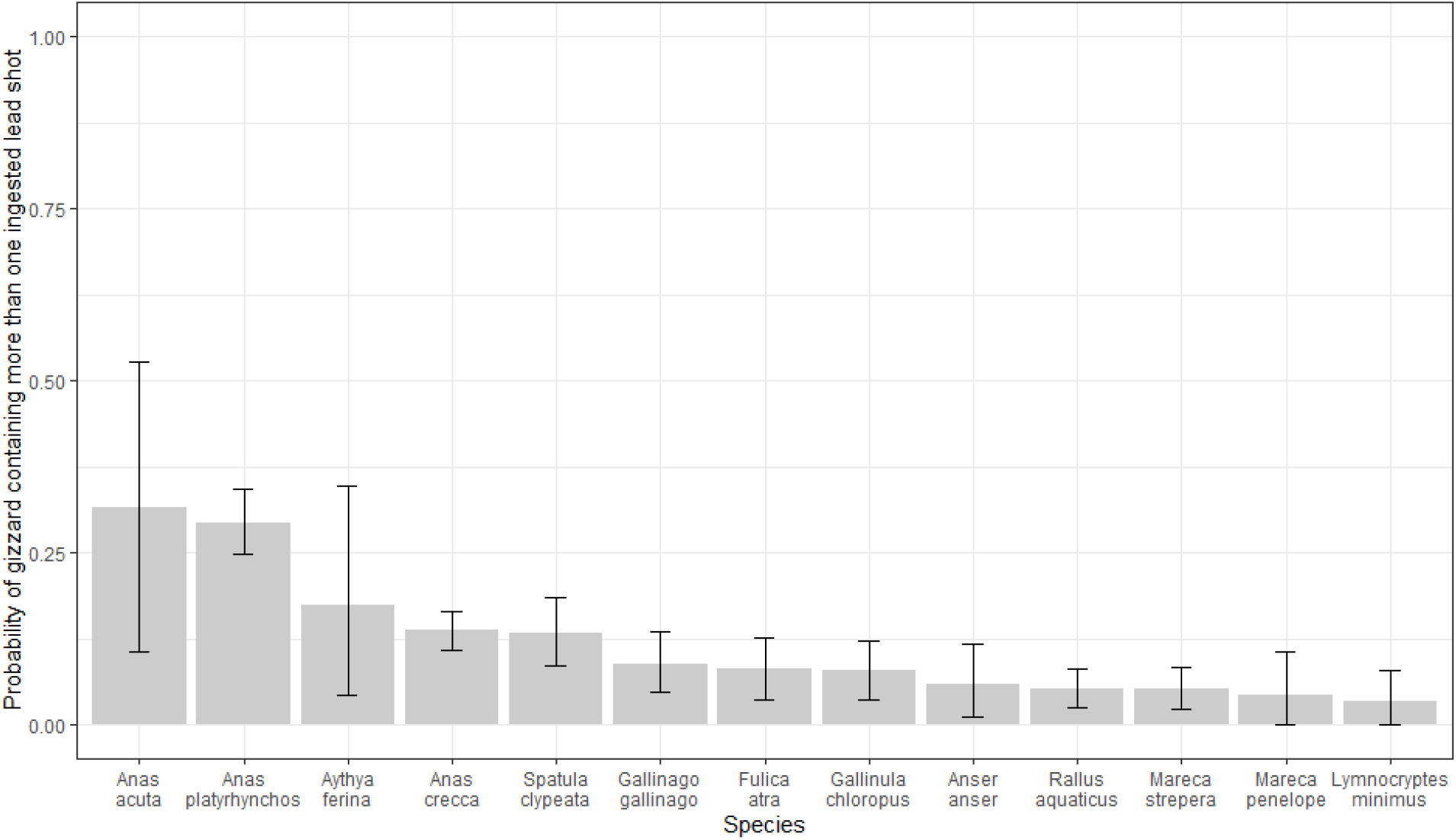
Prevalence of lead shot found in the gizzards of the 13 waterbird species hunted in the Camargue (south of France) from hunting season 1998/99 to 2017/18.

We did not detect any linear negative trend for lead shot prevalence in gizzard after winter 2007/2008 nor any linear decrease in lead shot numbers in gizzards with at least one lead (Table 3 and 4). Adding a smooth effect to assess for alternative time-trend shape of these variables did not improve the fit of the logistic model for prevalence (AICc = 3340 to compare to AICc of Table 3 and approximate significance of the smooth term *p* = 0.37). Similarly, such smooth effect did not significantly improve the fit of the model to the number of lead shots in gizzards (AICc = 5317 to compare to AICc of table 4 and *p* = 0.18). Again, only diet and family explained differences in lead shot prevalence in gizzards after the ban with a negative effect of both vegetarian diet and family Rallidae.

**Table 3.** Model selection for the variation of lead shot prevalence in the gizzard of the 13 species hunted in the Camargue after the ban from hunting season 2007/2008 onward to 2017/2018. Only models within 10 points of AIC with the best model are presented among the 8 models tested.

| Model | df | AICc | ΔAIC |
| --- | --- | --- | --- |
| Intercept + diet + family | 5 | 449.4 | 0 |
| Intercept + diet + family + year | 6 | 455.2 | 5.73 |
| Intercept + diet | 3 | 455.8 | 6.38 |
| Intercept+ family | 4 | 456.4 | 6.92 |
| Intercept | 2 | 458.6 | 9.15 |

**Table 4.** Model selection for annual variation in the number of lead shot found in the gizzards of the 13 species hunted in the Camargue after the ban from hunting season 2007/2008 onward to 2017/2018. Only models within 10 points of AIC with the best model are presented among the 8 models tested.

| **Model** | **df** | **AICc** | **ΔAICc** |
| --- | --- | --- | --- |
| Intercept + diet + family | 6 | 727.7 | 0 |
| Intercept + diet | 4 | 733.2 | 5.55 |
| Intercept + diet+ family + year | 7 | 733.3 | 5.59 |
| Intercept + family | 5 | 733.6 | 5.90 |
| Intercept | 3 | 735.9 | 8.17 |

Regarding nontoxic shots, model selection did not suggest an effect of lead ban on the prevalence of nontoxic shot, with no effect retained on the trend after model averaging (Table 5). Because of convergence issues due to low nontoxic shot abundance, we could not run a model with family and diet covariables to evaluate an increase in the number of nontoxic shot in gizzards from 2007 onwards. Yet, adding a year effect to a null intercept model was not retained as ΔAICc = 4.47 with the null model. Nevertheless, modelling an effect of year on the number of nontoxic shots in gizzards over the whole study period (1998-2017) showed a significant slight positive increase (β = 0.05 ± 0.02), which was however considered as a model equivalent to the null model since ΔAICc = 1.39). Adding a smooth effect on year did not improve the fit of the model.

**Table 5.** Model selection for the variation of the prevalence of nontoxic shots ingested in the gizzard of the 13 species hunted in the Camargue (1998-2017) before and after the ban on lead shot.

| **Model** | **df** | **AICc** | **ΔAICc** |
| --- | --- | --- | --- |
| Intercept + family | 4 | 707.0 | 0 |
| Intercept + family + ban | 5 | 707.8 | 0.76 |
| Intercept + diet | 3 | 743.8 | 36.8 |
| Intercept | 2 | 744.4 | 37.6 |
| Intercept + ban+ diet | 4 | 746.7 | 39.7 |
| Intercept + ban | 6 | 747.2 | 40.2 |

### Cartridge casings dataset

A total of 3966 cartridges were retrieved from the surrounding communal hunting trails of the Tour du Valat nature reserve. Markings were visible enough for all of them to be classified either as nontoxic or lead cartridges. The logistic model suggested a significant decrease in lead shot use by hunters across time (effect of year : β = -0.047 ± 0.0073, *p*<0.0001) from 2008 onwards, yet it remained high with more than half of cartridges found in the field being toxic lead cartridges even in 2018 and 2019 (Fig. 5).

**Figure 5.**
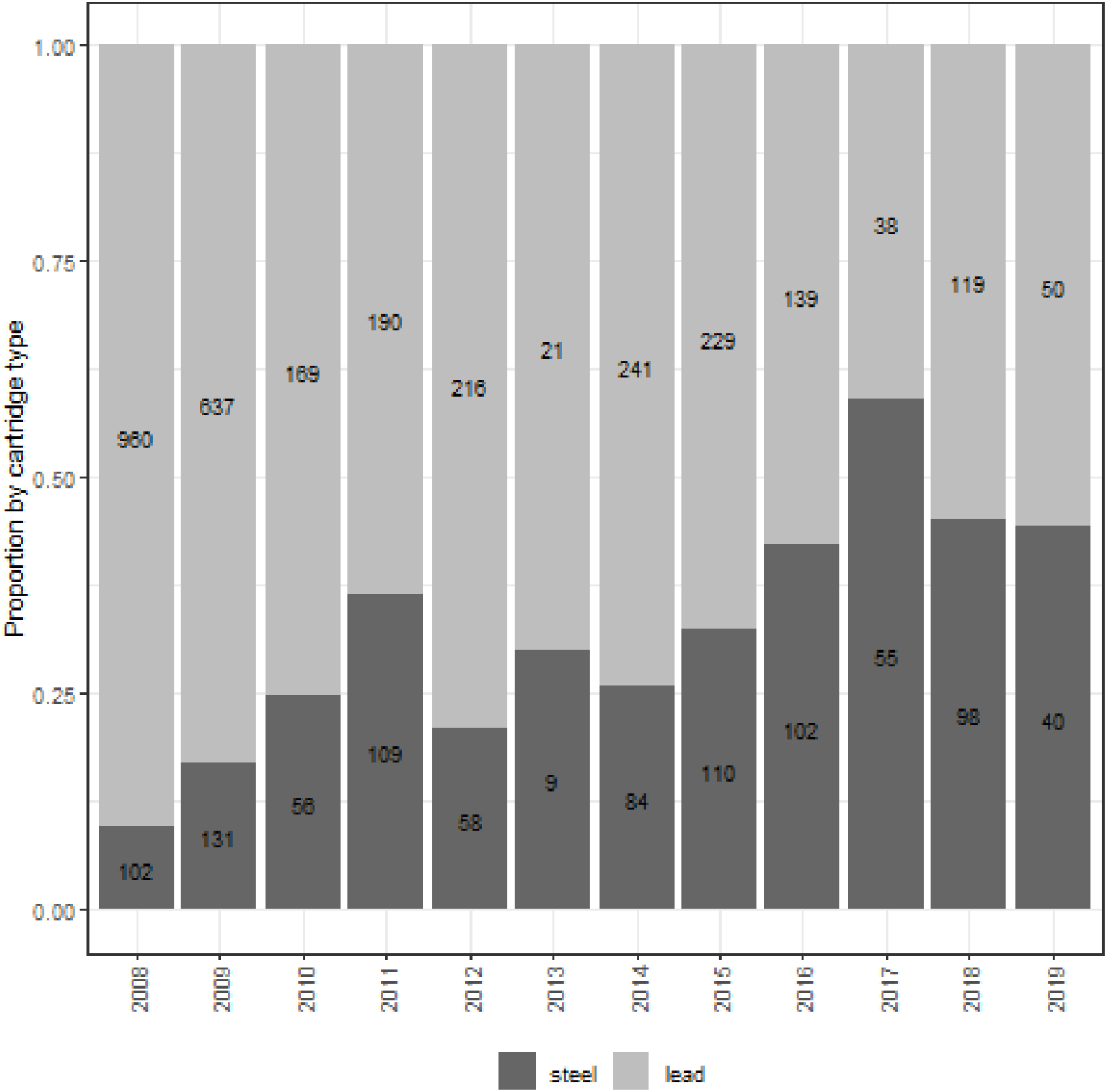
Proportion of cartridges made of lead or steel found at communal hunting trails surrounding Tour du Valat in the Camargue from hunting season 2008/2009 to 2019/2020. Values in the columns indicate the total number of cartridges of each type each year.

## Discussion

Despite a ten-year long ban on lead shot for waterbird hunting in France, our results do not show any significant reduction in their exposure to lead through shot ingestion in the Camargue. Indeed, gizzards of harvested waterfowl were as likely to contain at least one lead shot before than after the ban that occurred in 2006, with a mean prevalence of 12% over the 13 species considered across the study period. Since duck survival was shown to be negatively affected by lead from the first pellet found in the gizzard (Tavecchia et al. 2001), this result also indicates an absence of decrease in the expected demographic impact of lead contamination for wintering waterfowl in the south of France. It confirms, yet at a larger geographic scale and a longer time frame, the absence of effect of an earlier and localised lead ban at the scale of the sole Tour du Valat hunting group (Mondain-Monval et al. 2015). Our results are thus comparable to those from the UK, where lead shot ban has failed to reduce contamination over the subsequent 10 years, with an estimated 13 million ducks lethally poisoned since the implementation of the new regulation (Stroud et al. 2021). Conversely, our results contrast with a recent study in the Baltic sea where X-ray examination of Common eider *Somateria mollissima*, Velvet scoter *Melanitta fusca*, and Red-breasted merganser *Mergus serrator* could not find any imbedded or ingested pellet (Liljebäck et al. 2023). There, the remoteness of the breeding site located at the outer Archipelago of Oxelösund in the Baltic Sea, and the hunting restrictions on these species may explain the total absence of lead shot, either ingested or embedded.

In our case, the lack of decrease in contamination may stem from three different non-exclusive phenomena. First, the birds killed and analysed in this study do not only feed in the marshes of the Camargue and may have been contaminated elsewhere, in particular in countries without regulations on lead shot. Indeed, several long-distance migratory species (e.g. Eurasian teal *Anas crecca*) are likely to ingest lead while feeding during their migration across Russia and northern Europe. However, mallard *Anas platyrhynchos* (a similar granivorous species but mostly resident in the regions Guillemain et al. (2015)) do show a higher rate of lead prevalence (30%) than the migratory Eurasian teal (13%), suggesting that contamination is local to a large extent. Second, the local prevalence may result from an old contamination of wetlands, when lead was authorised. Indeed, it was shown that in the 1980s, Camargue wetland sediments presented one of the highest lead shot density worldwide with up to two million shots per hectares at some sites (Pain 1991). Given the long tradition of waterfowl hunting in the region (Tamisier and Dehorter 1999; Mondain-Monval et al. 2009), it is likely that most hunting marshes still contain high pre-ban lead shot concentration which remain available as grit for waterbirds even ten years after the ban. This would not be surprising since Kanstrup et al. (2020) showed that lead shot persisted in Danish wetland sediments more than 33 years after strict enforcement of the law prohibiting its use. Third, the contamination may also be enhanced by more recent local inputs of lead shots in wetlands, resulting from a lack of policy compliance regarding the lead ban. Indeed, the high proportion of lead cartridge casings found in the surrounding hunting spots of the nature reserve of Tour du Valat shows persistence of use of lead shot by hunters in wetlands despite a slow increase in steel shot use. This unequivocally indicates that the law is far from being strictly enforced in the Camargue. It is thus most likely that the absence of lead ban effectiveness results from a combination of the different phenomena described above.

Our results show large differences in lead shot contamination among waterfowl species (often more than ten times higher in particular for pintails, mallards and teals) in the Camargue than estimates compiled at the European level (Green and Pain 2016). In addition, for the five species of Anatidae for which we have sufficient data (*A. crecca, A. platyrhynchos, Mareca strepera, Mareca penelope, Spatula clypeata*), ingested lead prevalence is broadly comparable to previous results in the Camargue since the 1960s (Pirot 1978; Allouche 1983; Campredon 1984; Pirot and Taris 1987; Pain 1990), with the exception of the results of Pain (1990), who obtained much higher values for *A. platyrhynchos* (47%) and *S. clypeata* (23%).

Surprisingly, we also did not detect any increase in steel shot prevalence in gizzard after the ban on the use of lead while monitoring of cartridge casings show a progressive increase in the proportion of nontoxic ammunitions. Because nontoxic shots have been strictly used by the hunters of the Tour du Valat Estate since 1994 (Vallecillo et al. 2019), steel shots were already found in waterfowl gizzard before the French ban (Mondain-Monval et al. 2015). This may explain the lack of a before-after ban effect on this variable despite a slight long term increase.

Stroud et al. (2021) suggest that failure to decrease lead contamination in the UK resulted from the ban being partial, i.e. only applied to waterbird hunting. We concur with this analysis to explain the situation in France. Even in a country like France with a large and specialized body of civil servants (approximately 1700 environmental police staff) in charge of enforcing hunting/environmental laws, controls in the marshes are relatively rare. Furthermore, it is difficult to prove lead shot use by non-compliant hunters. Indeed, up to 2022/2023 hunting season, hunters had to be caught in the act with lead cartridges in the gun to be fined, while it was still perfectly legal to carry lead cartridges. In turn, when prosecution could be achieved, the relatively low amount of the fine in the event of such an offence (€135) does not really promote compliance. Finally, scepticism on lead toxicity, bad fame of nontoxic cartridges, which are more expensive and generally said to generate more crippling loss, remain common among hunters in the Camargue. This is despite numerous efforts made to raise awareness of local hunters, including blind-test clay-pigeon shooting sessions with nontoxic shots as well as numerous publications showing equivalent shooting effectiveness between lead and steel shot at the recommended maximum shooting distance of 30 meters (Mondain-Monval et al. 2015, 2017; Ellis and Miller 2022).

The new regulation (EC) no. 1907/2006 coming into force on this 2023/2024 hunting season within the European Union will likely contribute to solve these issues and facilitate law enforcement. Indeed it will not only prohibit the use of lead shot in wetlands but also the possession of lead cartridges by hunters in such habitats. This will certainly facilitate future controls by environmental police and gamekeepers.

While scientific efforts to reveal the effect of pollutants on wildlife often allows the implementation of new policies, the assessment of their effectiveness is rarely planned or funded. For instance, such assessment showed that the ineffectiveness of the enforcement of the policy to ban diclofenac from veterinary use in India because of its impact on vulture (Galligan et al. 2021). This work has allowed to propose several avenues for policy enhancement by helping to identify gaps in regulation effectiveness.

Regarding lead regulation, in order to be able to objectively judge the effectiveness of the new Regulation at European level and take into account the probable heterogeneity of the number of controls among each EU country, we strongly advocate initiating a coordinated international program to evaluate compliance to this new common regulation. We suggest that this scientific mid-term program, in cooperation with international hunting organisations, be based both on collections and analyses of a representative random sample of birds killed during hunting, as well as on a number of systematic hunters checks, in each country of the Union. Overall such a complete ban would help progressing toward a more sustainable practice of hunting in Europe (Kanstrup et al. 2018).

## Acknowledgements

We warmly thank all Tour du Valat and Camargue hunters, and in particular Jean-Pierre Reyre. We thank Gérard Panczer and Bernard Champagnon for the X-ray analyses carried out at the Centre commun de microspectrométries optiques CECOMO, Institut Lumière Matière, UMR5306, UCBL. We thank Elvin Miller, Marion Lourenço and Maël Olivier for helping with cartridge collection over the years. Finally, a great thank to Raquel Ambrosio for the preparation of the map of the study area. Finally, we acknowledge Matthieu Guillemain (OFB) for useful comments on a first draft of this paper.

---

**Appendix 1.** Species collected after harvest from hunting seasons 1998/1999 to 2017/2018 in the Camargue southern France. Diet, order, family, the number of gizzard collected (N), number of gizzard containing at least one lead N(lead) and at least one nontoxic shot N(nontoxic).

| Species | Diet | Order | Family | N | N (lead) | N (nontoxic) |
| --- | --- | --- | --- | --- | --- | --- |
| *Anas acuta* | omnivorous | Anseriformes | Anatidae | 19 | 6 | 2 |
| *Anas crecca* | omnivorous | Anseriformes | Anatidae | 546 | 75 | 23 |
| *Anas platyrhynchos* | omnivorous | Anseriformes | Anatidae | 337 | 102 | 30 |
| *Anas querquedula* | omnivorous | Anseriformes | Anatidae | 8 | 0 | 0 |
| *Anser anser* | vegetarian | Anseriformes | Anatidae | 86 | 5 | 15 |
| *Aythya ferina* | omnivorous | Anseriformes | Anatidae | 23 | 4 | 0 |
| *Aythya fuligula* | vegetarian | Anseriformes | Anatidae | 3 | 1 | 0 |
| *Calidris canutus* | carnivorous | Charadriiformes | Scolopacidae | 1 | 0 | 0 |
| *Coturnix coturnix* | granivorous | Galliformes | Phasianidae | 25 | 0 | 0 |
| *Fulica atra* | vegetarian | Gruiformes | Rallidae | 130 | 11 | 4 |
| *Gallinago gallinago* | carnivorous | Charadriiformes | Scolopacidae | 144 | 13 | 0 |
| *Gallinula chloropus* | omnivorous | Gruiformes | Rallidae | 150 | 11 | 6 |
| *Limosa limosa* | carnivorous | Charadriiformes | Scolopacidae | 6 | 0 | 0 |
| *Lymnocyptes minimus* | omnivorous | Charadriiformes | Scolopacidae | 86 | 3 | 0 |
| *Mareca penelope* | vegetarian | Anseriformes | Anatidae | 47 | 2 | 0 |
| *Mareca strepera* | vegetarian | Anseriformes | Anatidae | 214 | 10 | 8 |
| *Netta ruffina* | vegetarian | Anseriformes | Anatidae | 25 | 0 | 1 |
| *Numenius arquata* | carnivorous | Charadriiformes | Scolopacidae | 2 | 0 | 0 |
| *Numenius phaeopus* | carnivorous | Charadriiformes | Scolopacidae | 4 | 1 | 0 |
| *Phasianus colchicus* | granivorous | Galliformes | Phasianidae | 3 | 0 | 0 |
| *Pluvialis apricaria* | carnivorous | Charadriiformes | Scolopacidae | 4 | 0 | 0 |
| *Rallus aquaticus* | omnivorous | Gruiformes | Rallidae | 219 | 13 | 0 |
| *Scolopax rusticola* | omnivorous | Charadriiformes | Scolopacidae | 15 | 1 | 1 |
| *Spatula clypeata* | omnivorous | Anseriformes | Anatidae | 186 | 26 | 7 |
| *Streptopelia turtur* | vegetarian | Colombiformes | Columbidae | 5 | 0 | 0 |
| *Tringa nebularia* | carnivorous | Charadriiformes | Scolopacidae | 5 | 0 | 0 |
| *Tringa totanus* | carnivorous | Charadriiformes | Scolopacidae | 2 | 0 | 0 |
| *Vanellus vanellus* | carnivorous | Charadriiformes | Scolopacidae | 11 | 0 | 0 |

**Appendix 2.** Composition of alloy shots found in the gizzard of 14 birds harvested in the Camargue from 2004 to 2017. All gizzards containing alloy shots also contained at least one lead shot.

| # | species | date | N (lead) | N (steel) | N (alloy) | Alloy composition |
| --- | --- | --- | --- | --- | --- | --- |
| 1 | *Anas platyrhynchos* | 22/09/2004 | 1 | 0 | 1 | 78%Pb, 10%Zn, 6%W, 5%Sn |
| 2 | *Anas crecca* | 13/11/2004 | 4 | 0 | 1 | 56%Pb, 15%Si, 10%Al, 7%Sn |
| 3 | *Anas platyrhynchos* | 20/09/2006 | 5 | 0 | 1 | 80%Pb, 20% W |
| 4 | *Gallinula chloropus* | 04/11/2006 | 1 | 0 | 1 | 64%Pb, 20%Sn, 6%Al |
| 5 | *Spatula clypeata* | 24/12/2007 | 1 | 0 | 1 | 70%Pb, 20%Zn, 8%W |
| 6 | *Anas platyrhynchos* | 29/01/2008 | 1 | 0 | 1 | 68%Si, 23%Al, 6%Fe |
| 7 | *Anas crecca* | 23/12/2009 | 1 | 0 | 1 | 61%Bi, 23%Pb, 16%Sn |
| 8 | *Anas platyrhynchos* | 23/10/2010 | 4 | 0 | 1 | 46%Sn, 40%Pb, 6%W, 5%Si, 3%Al |
| 9 | *Anas crecca* | 16/12/2010 | 4 | 0 | 1 | 62%Pb, 26%Zn, 8%W |
| 10 | *Rallus aquaticus* | 12/10/2011 | 1 | 0 | 1 | 60%Pb, 40%Zn |
| 11 | *Anas platyrhynchos* | 01/11/2011 | 2 | 0 | 1 | 68%Pb, 25%W, 7%Zn |
| 12 | *Anas platyrhynchos* | 19/09/2012 | 17 | 0 | 1 | 77%Pb, 15%Sn |
| 13 | *Anser anser* | 29/11/2015 | 1 | 0 | 1 | 56% Cu, 17% W, 6% Fe |
| 14 | *Anas crecca* | 02/02/2017 | 4 | 0 | 1 | 60% Pb, 10% W |

